# A streamlined CRISPR workflow to introduce mutations and generate isogenic iPSCs for modeling amyotrophic lateral sclerosis

**DOI:** 10.1101/2021.06.21.449257

**Authors:** Eric Deneault, Mathilde Chaineau, Maria José Castellanos Montiel, Anna Kristyna Franco Flores, Ghazal Haghi, Carol X-Q Chen, Narges Abdian, Michael Nicouleau, Irina Shlaifer, Lenore K. Beitel, Thomas M. Durcan

**Affiliations:** The Neuro’s Early Drug Discovery Unit (EDDU), McGill University, 3801 University Street, Montreal, QC Canada H3A 2B4

**Keywords:** CRISPR, isogenic iPSC, ALS, *SOD1*-I114T, *SOD1*-G93A, *FUS*-H517Q

## Abstract

Amyotrophic lateral sclerosis (ALS) represents a complex neurodegenerative disorder with significant genetic heterogeneity. To date, both the genetic etiology and the underlying molecular mechanisms driving this disease remain poorly understood, although in recent years a number of studies have highlighted a number of genetic mutations causative for ALS. With these mutations pointing to potential pathways that may be affected within individuals with ALS, having the ability to generate human neurons and other disease relevant cells containing these mutations becomes even more critical if new therapies are to emerge. Recent developments with the advent of induced pluripotent stem cells (iPSCs) and clustered regularly interspaced short palindromic repeats (CRISPR) gene editing fields gave us the tools to introduce or correct a specific mutation at any site within the genome of an iPSC, and thus model the specific contribution of risk mutations. In this study we describe a rapid and efficient way to either introduce a mutation into a control line, or to correct a mutation, generating an isogenic control line from patient-derived iPSCs with a given mutation. The mutations introduced were the G93A mutation into *SOD1* or H517Q into *FUS*, and the mutation corrected was a patient iPSC line with I114T in *SOD1*. A combination of small molecules and growth factors were used to guide a stepwise differentiation of the edited cells into motor neurons in order to demonstrate that disease-relevant cells could be generated for downstream applications. Through a combination of iPSCs and CRISPR editing, the cells generated here will provide fundamental insights into the molecular mechanisms underlying neuron degeneration in ALS.

## 1. Introduction

Amyotrophic Lateral Sclerosis (ALS) is a fatal adult-onset neurodegenerative disorder characterized by a progressive loss of motor neurons (MNs). ALS patients have a median survival of 2-4 years [1]. Nearly 20% of ALS cases can be attributed to genetic variants identified in more than 20 ALS-associated genes involved primarily in RNA processing, protein homeostasis, oxidative stress and mitochondrial function [2]. The most frequent genes associated with ALS are *C9orf72, SOD1, TARDBP* and *FUS* [3]. *SOD1* encodes the detoxifying copper/zinc binding superoxide dismutase 1 enzyme, and most variants found are dominant missense mutations that cause similar protein structure variations and disease through gain-of-function mechanisms [4]. The most frequent *SOD1* mutations in North America and in the United Kingdom are A5V and I114T, respectively [5]. Nonetheless, I114T has demonstrated signs of reduced penetrance, emphasizing the genetic complexity of ALS [6]. Interestingly, the missense mutation G93A in human *SOD1* was shown to result in motor and sensory neuropathies in a transgenic mouse model that closely mimics ALS in humans [7]. *FUS* is a component of the heterogeneous nuclear ribonucleoprotein (hnRNP) complex involved in various RNA-associated processes, however FUS-positive aggregates observed in the cytosol do not include TDP-43, as observed in most ALS cases [8]. The missense mutation H517Q is the only autosomal recessive *FUS* mutation found associated with ALS [9]. However, the involvement of genetic variants in ALS pathology is difficult to prove due to the lack of effective models of human disease.

ALS modeling *in vitro* has accelerated since the advent of human iPSCs as these cells have the potential to generate disease-relevant cell types that can also be matched to the same genetic background as patients with the disease [10, 11]. A decade ago, the possibility of having almost unlimited access to human MNs from ALS patients without the need for invasive surgery became a reality following the development of protocols to guide iPSCs into spinal MNs [12]. These neurons now accelerate investigation into the pathophysiology of ALS. Importantly, the arrival of CRISPR genome editing provided a second state of the art technology, that when paired with iPSCs, meant we could correct or introduce any desired mutation, while also being able to knockout or ablate genes as required for functional studies [13]. CRISPR-Cas9 editing typically creates a break into the DNA at a targeted and sequence-specific genomic location. DNA repair mechanisms such as nonhomologous end-joining (NHEJ) or homology-directed repair (HDR) are then activated. The latter naturally happens less frequently but is more efficient for precise DNA modifications [14]. Precise HDR-based gene editing provides the means to knock-in disease-relevant point mutations or to correct such mutations back to the original sequence, creating an isogenic control line to pair with the patient iPSC. Merging both iPSC and CRISPR techniques facilitates the functional evaluation of ALS-associated variants across different genetic backgrounds. The penetrance of such variants can also be more easily evaluated using CRISPR editing, especially when no patient cell lines are available for reprogramming. Creating or correcting specific mutations in iPSCs to obtain isogenic controls, where the only difference is the target mutation, underlies direct causality between genotypes and phenotypes.

The methodologies presented here embody a specifically tailored strategy to knock-in or correct known genetic variants in the ALS genes *SOD1* (G93A and I114T) and *FUS* (H517Q) in human iPSCs derived from ALS patients or healthy individuals, combining CRISPR-Cas9 editing, droplet digital PCR (ddPCR) technology coupled with locked nucleic acid (LNA) probes, and antibiotic-free selection of modified cells. Following completion of our editing workflow, we further demonstrate that these edited cells can be efficiently differentiated into MNs, holding immense potential for developing models of ALS in a dish.

## 2. Materials and methods

**Table 1.** List of labware.

| Item | Supplier | Catalogue # |
| --- | --- | --- |
| 96-well clear flat bottom TC-treated culture microplate | ThermoFisher | 353072 |
| 384-well plate | ThermoFisher | 4309849 |
| Culture plate, 6-well | ThermoFisher | 087721B |
| ddPCR™96-well plates | BioRad | 12001925 |
| DG8 cartridge holder | BioRad | 1863051 |
| DG8 cartridges and gaskets | BioRad | 1864007 |
| Pierceable foil heat seal | BioRad | 1814040 |
| Pipet-Lite Multi Pipette L8-50XLS+ | RAININ Mettler Toledo | 17013804 |
| Tips GP-LTS-A-250 µL | RAININ Mettler Toledo | 30389278 |

Note: Please refer to the product datasheet from the supplier for further details on storage and preparation instructions.

**Table 2.** List of reagents.

| Item | Supplier | Catalogue # | Storage temp. |
| --- | --- | --- | --- |
| Accutase™ | StemCell Technologies | 07922 | –20°C |
| Alt-R® HDR Enhancer | Integrated DNA Technologies (IDT) | 1081073 | –20°C |
| Alt-R® S.p. HiFi Cas9 Nuclease V3 | IDT | 1081061 | –20°C |
| Amaxa™ P3 Primary Cell 4D Nucleofector™ X Kit S<br>- Buffer P3 Primary Cell Nucleofector™ Solution<br>- Supplement 1<br>- 16-well Nucleocuvette™ Strips | Lonza | V4XP-3032 | 4°C |
| Antibiotic-Antimycotic | Gibco | 15240-062 | Stock: –20°C<br>Working: 4°C |
| Antibody chicken anti-neurofilament heavy polypeptide antibody (NF-H) | Abcam | ab4680 | 4°C |
| Antibody goat anti choline-acetyltransferase (CHAT) | EMD Millipore | MAB144P | –20°C |
| Antibody goat anti-Sox1 | R&D System | AF3369 | –20°C |
| Antibody mouse anti homeobox 9 (HB9) | DSHB | 81.5C10-c | –20°C |
| Antibody mouse anti-ISLET1 (ISL1) | DSHB | 39.4D5-c | –20°C |
| Antibody mouse anti-Ki67 | BD Biosciences | 556003 | 4°C |
| Antibody rabbit anti-Nestin | Abcam | ab92391 | –20°C |
| Aqua-Poly/Mount | Polysciences | 18606-5 | RT |
| Ascorbic acid | Sigma-Aldrich | A5960 | Stock: –20°C<br>Working: 4°C |
| B27 | Life Technologies | 17504-044 | Stock: –20°C<br>Working: 4°C |
| Bovine serum albumin (BSA) | Multicell | 800-095-CG | 4°C |
| Brain-derived neurotrophic factor (BDNF) | Peprotech | 450-02 | Stock: –20°C<br>Working: 4°C |
| CHIR99021 | Selleckchem | S2924 | Stock: –20°C<br>Working: 4°C |
| Ciliary neurotrophic factor | Peprotech | 450-13 | Stock: –20°C<br>Working: 4°C |
| Compound E | StemCell Technologies | 73954 | Stock: –20°C<br>Working: 4°C |
| ddPCR Supermix for Probes (no dUTP) | BioRad | 1863024 | –20°C |
| DMEM/F12 | Gibco | 10565-018 | 4°C |
| DMH1 | Selleckchem | S7146 | Stock: –20°C<br>Working: 4°C |
| DMSO | Fisher Scientific | BP231-1 | RT |
| dNTPs | New England BioLabs | N0447L | –20°C |
| Dithiothreitol (DTT) | ThermoFisher | NP0004 | –20°C |
| Droplet Generation Oil for Probes | BioRad | 1864110 | RT |
| Droplet Reader Oil | BioRad | 1863004 | RT |
| Fetal Bovine Serum (FBS) | Life Technologies | 12484-028 | Stock: –20°C<br>Working: 4°C |
| First strand buffer | Teknova | F1204 | –20°C |
| Formaldehyde | ThermoFisher | 28908 | RT |
| Gentle Cell Dissociation Reagent | StemCell Technologies | 07174 | RT |
| Hoechst 33342 | Life Technologies | H3570 | 4°C |
| Insulin like growth factor-1 | Peprtech | 100-11 | Stock: –20°C<br>Working: 4°C |
| Laminin | Sigma-Aldrich | L2020 | Stock: –20°C<br>Working: 4°C |
| LNA™ Probes | IDT | N/A (custom order) | –20°C<sup>†</sup> |
| Matrigel Matrix hESC-qualified | Corning Millipore | 354277 | Stock: –80°C<br>Working: 4°C* |
| miRNeasy Micro Kit | Qiagen | 217004 | RT |
| M-MLV RT | ThermoFisher | 28025013 | –20°C |
| mTeSR1™ | StemCell Technologies | 85850 | Stock: –20°C<br>Working: 4°C |
| N2 | Life Technologies | 17502-048 | Stock: –20°C<br>Working: 4°C |
| Neurobasal medium | Life Technologies | 21103-049 | 4°C |
| Normal Donkey Serum | EMD Millipore | S30-100 | Stock: –20°C<br>Working: 4°C |
| PCR cloning kit | New England Biolabs | E1202S | –20°C |
| PCR primers | Invitrogen | N/A (custom order) | –20°C |
| Phosphate-Buffered Saline (PBS) | ThermoFisher | 10010023 | RT |
| Poly-L-Ornithine (PLO) | Sigma-Aldrich | P3655 | –20°C |
| Purmorphamine | Sigma-Aldrich | SML-0868 | Stock: –20°C<br>Working: 4°C |
| qPCR assay - ACTB | ThermoFisher | Hs01060665_g1 | –20°C |
| qPCR assay - ChAT | ThermoFisher | Hs00758143_m1 | –20°C |
| qPCR assay - GAPDH | ThermoFisher | Hs02786624_g1 | –20°C |
| qPCR assay – Hb9 | ThermoFisher | Hs00907365_m1 | –20°C |
| qPCR assay – Islet-1 | ThermoFisher | Hs00158126_m1 | –20°C |
| qPCR assay – MAP2 | ThermoFisher | Hs00258900_m1 | –20°C |
| qPCR assay - NANOG | ThermoFisher | Hs02387400_g1 | –20°C |
| qPCR assay – OLIG2 | ThermoFisher | Hs00377820_m1 | –20°C |
| qPCR assay – Pax6 | ThermoFisher | Hs01088114_m1 | –20°C |
| qPCR assay – POU5F1 | ThermoFisher | Hs04260367_gH | –20°C |
| QuickExtract™ DNA Extraction Solution | Lucigen | QE09050 | –20°C |
| Random primers | ThermoFisher | 48190011 | –20°C |
| ReLeSR™ | StemCell Technologies | 05872 | RT |
| Retinoic acid (RA) | Sigma-Aldrich | R2625 | Stock: –20°C<br>Working: 4°C |
| SB431542 | Selleckchem | S1067 | Stock: –20°C<br>Working: 4°C |
| Secondary antibodies raised in various species and conjugated to DyLight 488, 550 or Alexa Fluor 647 | Abcam or Jackson ImmunoResearch | N/A | 4°C |
| sgRNA | Synthego | N/A (custom order) | –20°C |
| Taqman™ Fast Advanced Master Mix, no UNG | Applied Biosystems | A44360 | –20°C |
| Triton™ X-100 | BioShop Canada | TRX777.500 | RT |
| Ultramer® ssODN | IDT | N/A (custom order) | –20°C |
| UltraPure™ DNase/RNase-Free Distilled Water (H<sub>2</sub>O) | ThermoFisher | 10977015 | RT |
| Y-27632 (ROCK inhibitor) | Selleckchem | S1049 | Stock: –80°C<br>Working: 4°C |

\*Matrigel working solution must be used immediately or stored at –20°C for later use; †Protect from light; N/A = not available

**Table 3.** List of equipment.

| Item | Supplier | Catalogue # |
| --- | --- | --- |
| Biosafety Cabinet | Baker | SG404P |
| C1000 Touch™ thermal cycler with 96–deep well reaction module | BioRad | 1851197 |
| Cell Counter | Logos Biosystems | L40002 |
| Cell culture incubator | ThermoScientific | Steri-Cycle Model 370 |
| Centrifuge | Eppendorf | 022626001 |
| Heat Block | VWR | 12621-088 |
| Microscope | Motic | 110010380054 |
| Nucleofector 4D™ Core Unit | Lonza | AAF-1002B |
| PCR Plate Spinner | VWR | 89184-608 |
| PX1™ PCR Plate Sealer | BioRad | 1814000 |
| QuantStudio 5 | ThermoFisher | A28140 |
| QX200™ Droplet Generator | BioRad | 1864002 |
| QX200™ Droplet Reader | BioRad | 1864003 |
| Vortex Mixer | Fisher Scientific | 88880017 |

### 2.1. Cell lines and iPSC reprogramming

The use of stem cells in this research was approved by the McGill University Health Centre Research Ethics Board (DURCAN_IPSC/ 2019-5374). The iPSC line TD-15 (*SOD1*^I114T/+^) was reprogrammed from a 45-year-old Caucasian female with ALS and carrying the heterozygous mutation c.341T>C, p.I114T in the *SOD1* gene. To knock-in selected mutations, we used the noncarrier AIW002-02 iPSC line, reprogrammed from a 37-year-old Caucasian male as previously described [15]. Briefly, peripheral blood mononuclear cells (PBMCs) were collected for reprogramming all iPSC lines. Different approaches were used to overexpress *OCT4/POU5F1, SOX2, KLF4* and *MYC* for cell reprogramming, i.e., Sendai virus for AIW002-02, and episomal for *SOD1*^I114T/+^. Emerging iPSC colonies were selected for activated endogenous human pluripotency markers, differentiation potential into three germ layer cells after embryoid body formation *in vitro*, and normal karyotype. Short tandem repeat (STR) analysis was performed at The Centre for Applied Genomics (TCAG, Canada).

### 2.2. Design of single guide RNAs

The design of our single guide RNAs (sgRNAs) was made using tools available at www.benchling.com. In brief, ~50-75 base pairs (bp) of genomic DNA sequence on each side of the mutation to edit were used to search for the best-scored sgRNA. We considered a sgRNA to be good if:

1. It cut less than 10 bp away from the mutation to edit
2. It had a low off-target event potential (= highest off-target score in www.benchling.com) We were mindful that some single nucleotide polymorphisms (SNPs) present in the genome of the patient iPSC line, but not in the reference genome in which the design was made, or might create new mismatches in the sgRNA sequence, and vice versa, so we tried to avoid any SNP within the chosen sgRNA sequence through using tools such as www.ensembl.org.

### 2.3. Design of single-stranded oligodeoxynucleotide

For HDR-based gene editing, we typically designed a DNA template with 40-60 bp homology arms flanking the appropriate codon to change. This template was synthesized as single-stranded oligodeoxynucleotide (ssODN). We sought to disrupt the proto-adjacent motif (PAM) of the sgRNA sequence embedded within the ssODN sequence by introducing a silent mutation(s) when possible to prevent Cas9 from cutting the template and the newly engineered allele. Silent mutations were introduced by changing one of the two Gs within the PAM of the sgRNA with another nucleotide resulting in no amino acid change after translation, or by changing other nucleotides close to the PAM sequence. When introducing such silent mutations, we payed close attention to the human codon usage frequency as some codons are rarely used in humans, and tried to not create new donor or acceptor splicing sites, i.e., new GTs or AGs, that might disrupt the expected splicing pattern.

### 2.4. Design of primers and probes

The detection of the modified nucleotide by ddPCR was based on a TaqMan^®^ assay including two PCR primers and two DNA probes fused with different fluorophores (FAM and HEX). One probe was specific to the original allele and the other probe to the edited allele. Locked Nucleic Acid (LNA^®^) probes were designed following the manufacturer criteria outlined below:

1. The SNP of interest should be positioned in the center of the probe
2. The difference between melting temperature (T_m_) of match and T_m_ of mismatch hybridizations (ΔT_m_) should be 8°C or more. ΔT_m_ is of particular importance in applications where probes are used for specific detection of mutations, or homologous sequences. A large ΔT_m_ is required to ensure specific detection of the sequence of interest. As an example, if the wildtype allele is an A, and the mutant allele is a G, mismatches would be A:C and G:T. Use a thermodynamics calculator (www.idtdna.com/calc/analyzer/lna) to calculate match and mismatch T_m_’s of possible probe configurations. Enter the Mg^+2^ and dNTP concentrations for qPCR conditions (3 mM MgCl2 and 0.8 mM dNTPs)
3. Enter the probe target sequence with LNA bases denoted with a “+” in front of the base of interest (+A, +C, +G, +T). Start with an LNA base on the SNP, and one on each adjacent base. Match Tm: 64-68°C. Mismatch T_m_ no higher than 56°C (again, minimum ΔT_m_ of 8°C, greater is better). It may be necessary to design towards the antisense strand if T_m_ mismatch is troublesome for the sense strand. If this is the case, make sure to design both probes to the same strand so they are non-complementary
4. Additional LNA bases are added to achieve optimal T_m_ mismatch. Some mismatches can make achieving optimal thermodynamics challenging. Examples include designs involving G-T and C-A mismatches. Ideal configuration:

- LNA on SNP and on adjacent bases (3 LNAs in a row), e.g., ACGT+A+C+TATCG.
- Do not place more than 4 LNAs in a row. No more than 6 LNAs per probe.
- Spread out the LNAs to get a greater effect on T_m_
- Avoid stretches of 3+ C’s or G’s
- Placing LNAs on G and C will give the greatest T_m_ increase
- No LNA on first base. Avoid LNA on second base
- Avoid probes with a 5’ G residue (can quench fluorophore)
- Shorter LNA probes (10-14 nt) are more effective at single base discrimination (ensure T_m_ ≥ 64°C)
5. Use PrimerQuest or PrimerBlast to design a set of primers surrounding SNP of interest: Primer T_m_: Aim for 62°C (60-63°C). Amplicon size ~80-150 bp
6. Check primers for dimers, hairpins, heterodimers using OligoAnalyzer (https://www.idtdna.com/calc/analyzer). BLAST primers/probe to check for specificity/off target transcripts

### 2.5. Preparation of ribonucleoprotein complex

The CRISPR components [Cas9 + sgRNA + ssODN] were mixed as follows: 1 μl of Cas9 protein (stock 61 μM) with 3 μl of sgRNA (stock 100 μM) at room temperature (RT) for 10-20 minutes. This mix forms a Cas9:sgRNA ribonucleoprotein (RNP) complex. After formation of the RNP complex, add 1 μl of ssODN (stock 100 μM) and 20 μl of nucleofection buffer P3 to the RNP mix.

### 2.6. iPSC nucleofection

When iPSCs attain a maximum of 50% confluency:

1. Wash iPSCs with PBS
2. Detach with Accutase at 37°C for 10 minutes
3. Centrifuge 500,000 detached cells in a separate tube at 250g for 3 minutes
4. Wash with PBS
5. Centrifuge
6. Remove supernatant
7. Resuspend the pellet gently with the 25 μl of the RNP-ssODN-buffer mix
8. Transfer the resuspension into a 16-strip cuvette
9. Nucleofect using the CA137program in a Nucleofector 4D device
10. Add 75 μl of [mTeSR + ROCK inhibitor 10 μM final] on top of the cells in the cuvette
11. Transfer the content of the cuvette into a new 15-ml conical tube containing 10 ml of nucleofection media [mTeSR + ROCK inhibitor 10 μM final + HDR enhancer 30 μM final]
12. Distribute 100 μl of this mix per well in a Matrigel-coated 96-well plate, after aspirating the Matrigel
13. Replace the culture media with 100 μl per well of iPSC maintenance media [mTeSR + ROCK inhibitor 10 μM final] after 24 hours, and then daily with mTeSR after 48 hours

### 2.7. Quantification of edited cells

When 80-90% confluent, i.e., 10-14 days post-nucleofection:

1. Wash cells twice with 100 μl/well of PBS
2. Treat with 30 μl/well of Accutase for 10 minutes at 37°C
3. Transfer half of each well (15 μl) into a new Matrigel-coated 96-well plate pre-filled with 150 μl/well of mTeSR + ROCK inhibitor 10 μM
4. Store the backup plate in the incubator at 37°C
5. Add 35 μl of QuickExtract solution to the remaining cells/Accutase mix in the original plate
6. Heat at 65°C for 10 minutes, then 95°C for 5 minutes
7. Dilute the DNA extract 1/20 with H_2_O to evaluate the proportion of edited alleles using ddPCR
8. Prepare a typical ddPCR master mix for a whole 96-well plate by mixing 1,170 μl of “ddPCR Supermix for Probes (no dUTP)”, 975 μl H_2_O, 20.8 μl of each primer (stock 100 μM) and 6.5 μl of each probe (stock 100 μM)
9. Distribute 19 μl per well of the master mix into a new 96-well ddPCR plate
10. Add 2 μl per well of the diluted DNA extract into each corresponding well The droplets were assembled using the QX200™ Droplet Generator according to the manufacturer’s protocol, and transferred into a ddPCR™96-well plate. The PCR reaction was performed using the C1000 Touch™ thermal cycler with 96–deep well reaction module. The resulting droplets were analysed using the QX200™ Droplet Reader, and the absolute quantification of edited and unedited alleles was obtained.

### 2.8. Passage at limiting dilution

If no well was identified with 100% edited alleles, we searched for the well(s) with the highest proportion of edited alleles. When the corresponding well in the backup plate from the previous section reach 60-70% confluency:

1. Wash cells twice with 100 μl PBS per well
2. Treat with 30 μl per well of Accutase at 37°C until all cells lift off
3. Add 250 μl of mTeSR + ROCK inhibitor 10 μM per well and mix gently by pipetting up and down 2-3 times
4. Count cells
5. Transfer 100-200 cells into a new 15-ml conical tube previously filled with 10 ml of mTeSR + ROCK inhibitor 10 μM
6. Distribute 100 μl/well of this mix in a new Matrigel-coated 96-well plate Keep in mind that the number of cells to transfer can differ considerably between different iPSC lines, depending on robustness, survival, growth rate, etc. This can be assessed beforehand with the goal of ending up with ~1 colony per well. This passaging/enrichment step was repeated until a well with 100% edited cells was found.

### 2.9. Seguence Integrity

Isolated cell populations were assessed for sequence integrity and homozygosity/heterozygosity of the mutation by PCR cloning and Sanger sequencing. We designed PCR primers that gave rise to a ~300 bp single/clean band on an agarose gel with the mutation of interest located in the middle. This fragment was used for PCR cloning or directly for Sanger sequencing using the same PCR primers. We also monitored genomic integrity by karyotyping and/or targeted PCR.

### 2.10. Generation of spinal MNs from human iPSCs

Human AIW002-02 and *SOD1*^G93A/G93A^ iPSC lines were differentiated into human motor neurons progenitor cells (MN NPCs) and finally into motor neurons (MNs) according to a previously published protocol, with some modifications outlined below [16]:

1. Dissociate iPSCs using Gentle Cell Dissociation Reagent and split at a 1:5 ratio into Matrigel-coated T25 flasks, in a neural medium composed of DMEM/F12 and Neurobasal medium at a 1:1 ratio, 0.5x B27 and 0.5x N2 and 1x antibiotic-antimycotic and 100 μM ascorbic acid (AA), enriched with 3 μM CHIR99021, 2 μM SB431542 and 2 μM DMH1, and fully changed every other day for 6 days
2. For the first 24 hours, supplement the media with 10 μM ROCK inhibitor Y-27632
3. After 6 days, split the cells into 10 μg/ml Poly-L-ornithine (PLO) and 5μg/ml laminin coated flasks at a ratio of 1:3 to 1:6 with Gentle Cell Dissociation Reagent. This is in the same neural medium described previously, supplemented with 1 μM CHIR99021, 2 μM SB431542, 2 μM DMH1, 0.1 μM retinoic acid (RA) and 0.5 μM Purmorphamine, enriched with ROCK inhibitor Y-27632 for the first 24 hours. Change the media every other day
4. After 6 days (day 12), split the cells into 10 μg/ml Poly-L-ornithine (PLO) and 5μg/ml laminin coated flasks at a ratio 1:3 to 1:6 with Gentle Cell Dissociation Reagent, in the same neural medium described previously, supplemented with 3 μM CHIR99021, 2 μM SB431542, 2 μM DMH1, 0.1 μM RA and 0.5 μM Purmorphamine, and 0.5 mM VPA, enriched with ROCK inhibitor Y-27632 for the first 24 hours
5. Split and culture the MNPCs in the same media for subsequent differentiation into MNs or freeze down in a freezing medium composed of 10% DMSO in Fetal Bovine Serum (FBS) in order to constitute a stock of MNPCs
6. For differentiation into MNs, dissociate MNPCs with Accutase and plate onto PLO and Laminin coated coverslips at a density of 50,000 cells per coverslip into the neural medium described previously supplemented with 0.5 μM RA, 0.1 μM Purmorphamine, 0.1 μM Compound E, and 10 ng/ml insulin like growth factor-1, brain-derived neurotrophic factor and ciliary neurotrophic factor, with ROCK inhibitor Y-27632 added for the first 24 hours
7. Differentiate MNs for 2 or 4 weeks and perform half-changes of the differentiation media once a week

### 2.11. Characterization of human iPSCs-derived MNs by immunocytochemistry

AIW002-02 and *SOD1*^G93A/G93A^ iPSCs-derived MN NPCs were characterized by immunocytochemistry using the following primary antibodies: goat anti-Sox1 (1/100), rabbit anti-Nestin (1/250) and mouse anti-Ki67 (1/50). AIW002-02 and *SOD1*^G93A/G93A^ iPSC-derived MNs were characterized by immunocytochemistry after 28 days of differentiation using the following primary antibodies: mouse anti homeobox 9 (HB9), mouse anti-ISLET1 (ISL1), goat anti choline-acetyltransferase (CHAT) and chicken anti-neurofilament heavy polypeptide antibody (NF-H). Secondary antibodies against primary antibodies raised in various species were conjugated to DyLight 488, 550 or Alexa Fluor 647 (all at 1/500). Briefly:

1. Fix cells on coverslips with 4% formaldehyde in PBS for 15 minutes
2. Wash with PBS
3. Permeabilize with 0.2% Triton X-100 in PBS for 10 minutes
4. Block in a blocking solution composed of 5% normal donkey serum, 1% bovine serum albumin (BSA) and 0.02% Triton X-100 in PBS for 45 minutes
5. Incubate cells overnight at 4°C with primary antibodies described previously that were diluted in the blocking solution
6. Wash 3 times in PBS
7. Incubate 1 hour at RT in the dark with secondary antibodies diluted in the blocking solution
8. Wash cells 2 times in PBS
9. Incubate for 10 minutes with Hoechst 33342 (1/5000 in PBS)
10. Wash once in PBS
11. Mount coverslips with Aqua-Poly/Mount and acquire images using a confocal microscope

### 2.12. qPCR analysis of iPSCs, MNPCs and MNs

Total RNAs were isolated using the miRNeasy Micro Kit according to the manufacturer’s instructions:

1. Reverse transcription was done on the total RNA fraction in order to obtain cDNA in a total volume of 40 μl containing 100 ng total RNA, 12.5 ng/μl random primers, 0.5 mM dNTPs, 0.01 M DTT, 1x first strand buffer and 400 U M-MLV RT
2. Each total cDNA mix was diluted 1:4 after reverse transcription
3. The qPCR reactions were performed in a total volume of 10 μl on a 384-well plate using the QuantStudio 5 PCR machine
4. For each well, PCR mix included 9 μl of 2X Taqman™ Fast Advanced Master Mix no UNG, 0.5 μl primers/probe mix, 1 μl cDNA, with H_2_O up to 10 μl
5. Serial dilutions of a mix of cDNAs ranging between 0.003052 ng and 50 ng were used to generate a calibration curve for absolute quantification

Expression levels were given as a ratio between the relative quantities of the gene of interest and the endogenous controls. The mean between β-actin and GAPDH was used as endogenous controls for normalization.

## 3. Results

### 3.1. Nucleotide identification

In this study, we present a streamlined approach to CRISPR-Cas9 gene editing, with a focus on *SOD1* and *FUS* mutations. The key steps of our method are illustrated in **Figure 1A.** In order to correct specific mutations in carrier cells, or conversely to knock-in selected mutations into non-carrier cells, we first sought to locate the right nucleotides using the human genome browser Ensembl (**Fig. 1A**). We acknowledge that the mutation ID may be confusing if it has been assigned according to the original transcripts discovered for its corresponding gene. For instance, regarding the missense mutation I114T in *SOD1*, we searched for an isoleucine (I) as the 114th amino acid on a protein-coding transcript corresponding to the gene *SOD1*. However, there are actually two known protein-coding transcripts corresponding to *SOD1*, and only one of these transcripts has an isoleucine at position 114, i.e., transcript SOD1-201 (ENST00000270142.10 in http://useast.ensembl.org/). In this case, the data regarding the SNP rs121912441 in dbSNP (https://www.ncbi.nlm.nih.gov/snp/) was essential to ensure that we targeted the right nucleotide. We applied this same approach for the precise localization of the other codons to substitute, i.e., G93A in *SOD1*, as well as H517Q in *FUS* (**Table 4**). Thus, with our methodologies in place, we focused on correcting the *SOD1*-I114T missense mutation in the ALS patient iPSC line TD-15 (*SOD1*^I114T/+^), and to knock-in the *SOD1*-G93A and FUS-H517Q mutations into our control iPSC line AIW002-02 (**Table 4**). For each target mutation, we designed a sgRNA and a single-stranded oligonucleotide (ssODN) template that were used for homology-directed repair (HDR)-based nucleotide substitution using CRISPR-Cas9 gene editing. Each sgRNA was designed to promote a DNA break as close as possible to the target codon (nucleotides in green in **Table 4**). The nucleotide substitution creating the mutation G93A in *SOD1* also disrupted the PAM sequence such that Cas9 was prevented from cutting the newly engineered allele. Nucleotide substitution for *SOD1*-I114T or *FUS*-H517Q did not disrupt the PAM sequence in the corresponding ssODN sequence but was relatively close (one and four bp away, respectively), hence no additional silent mutations were introduced.

**Fig. 1.**
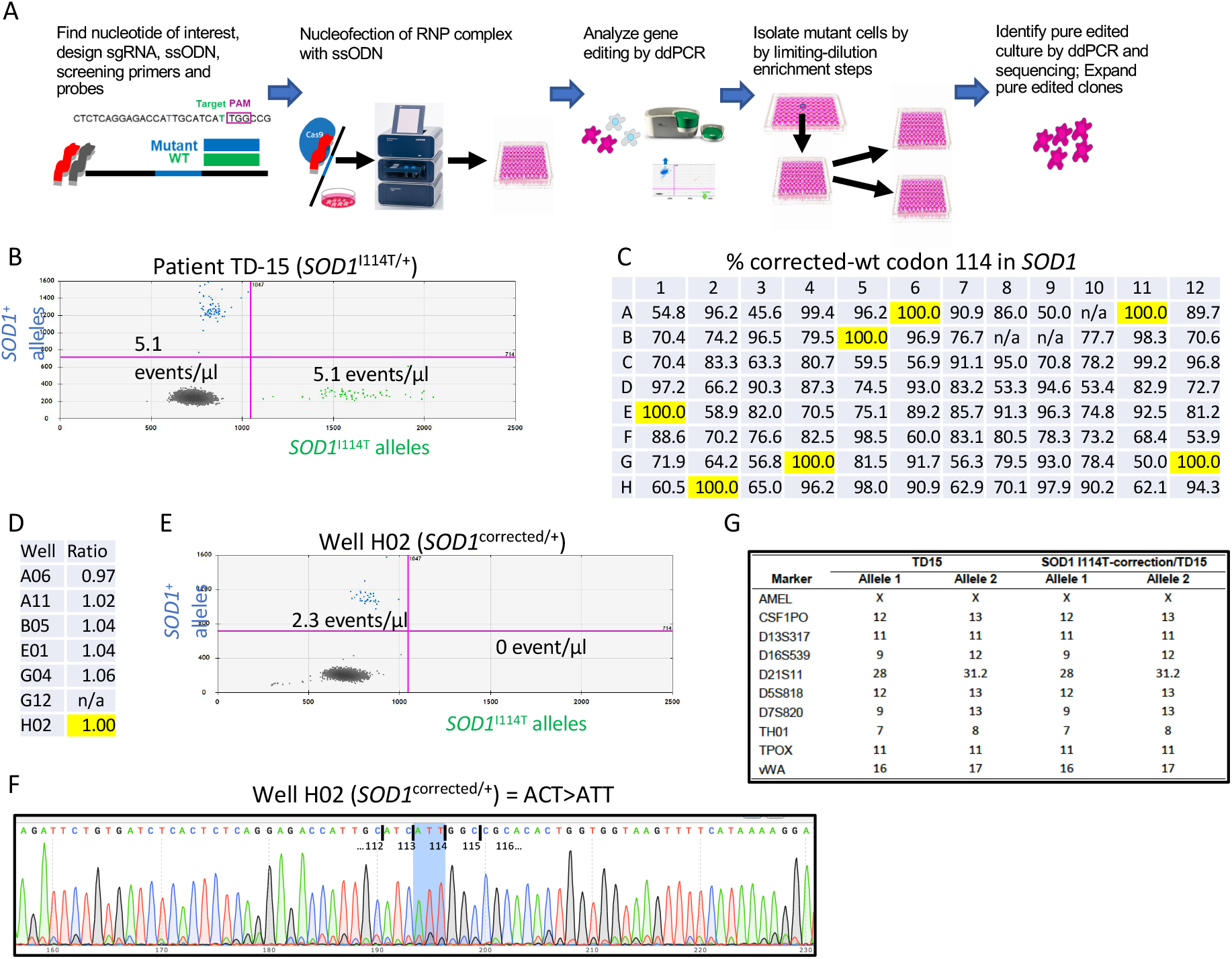
Correction of the I114T mutation in *SOD1* in the patient-derived iPSC line *SOD1*^I114T/+^. (A) Gene editing strategy including selection of target nucleotide considering SNPs (as per https://www.ncbi.nlm.nih.gov/snp/), design of sgRNAs via Cloud-Based Informatics Platform for Life Sciences R&D (Benchling), design of ssODN to contain target mutation with homology arms, design of LNA probes with distinct fluorophores (using tools available on https://www.idtdna.com/), and primers using Primer designing tool (nih.gov). Cells are nucleofected using the Nucleofector 4D™ System with RNP complex (Cas9+gRNA) +ssODN and cultured on 96 well plate. The resulting cell colonies are screened for editing event by ddPCR (QX200™ Droplet PCR System, Bio-Rad) using probes to quantify edited vs unedited alleles. The well with the highest % of edited cells is then selected for subsequent enrichment by limiting-dilution to isolate the edited clones. Pure edited cells are confirmed by ddPCR and Sanger sequencing. (B) Scatter plot showing the distribution of wt (blue) vs mutant (green) alleles in the heterozygous patient’s iPSC line *SOD1*^I114T/+^. (C) Percentages of corrected (or wt) alleles in each well of a 96-well plate measured by ddPCR in the absolute quantification mode. Wells selected for zygosity assessment are highlighted in yellow. (D) Zygosity of the corrected (or wt) codon 114 in iPSCs from wells selected in (C), presented as the ratio of *SOD1*^I114T/+^:*USP19*^+^ alleles. (E) Scatter plot showing the distribution of wt (blue) vs mutant (green) alleles in iPSCs from well H02 selected in (C). (F) Sanger sequence chromatogram from iPSCs in the selected well H02; shaded blue area shows complete correction of target codon. (G) Short tandem repeat analysis of line *SOD1*^I114T/+^ and cells from well H02. n/a = no colony grew in this well.

**Table 4.**
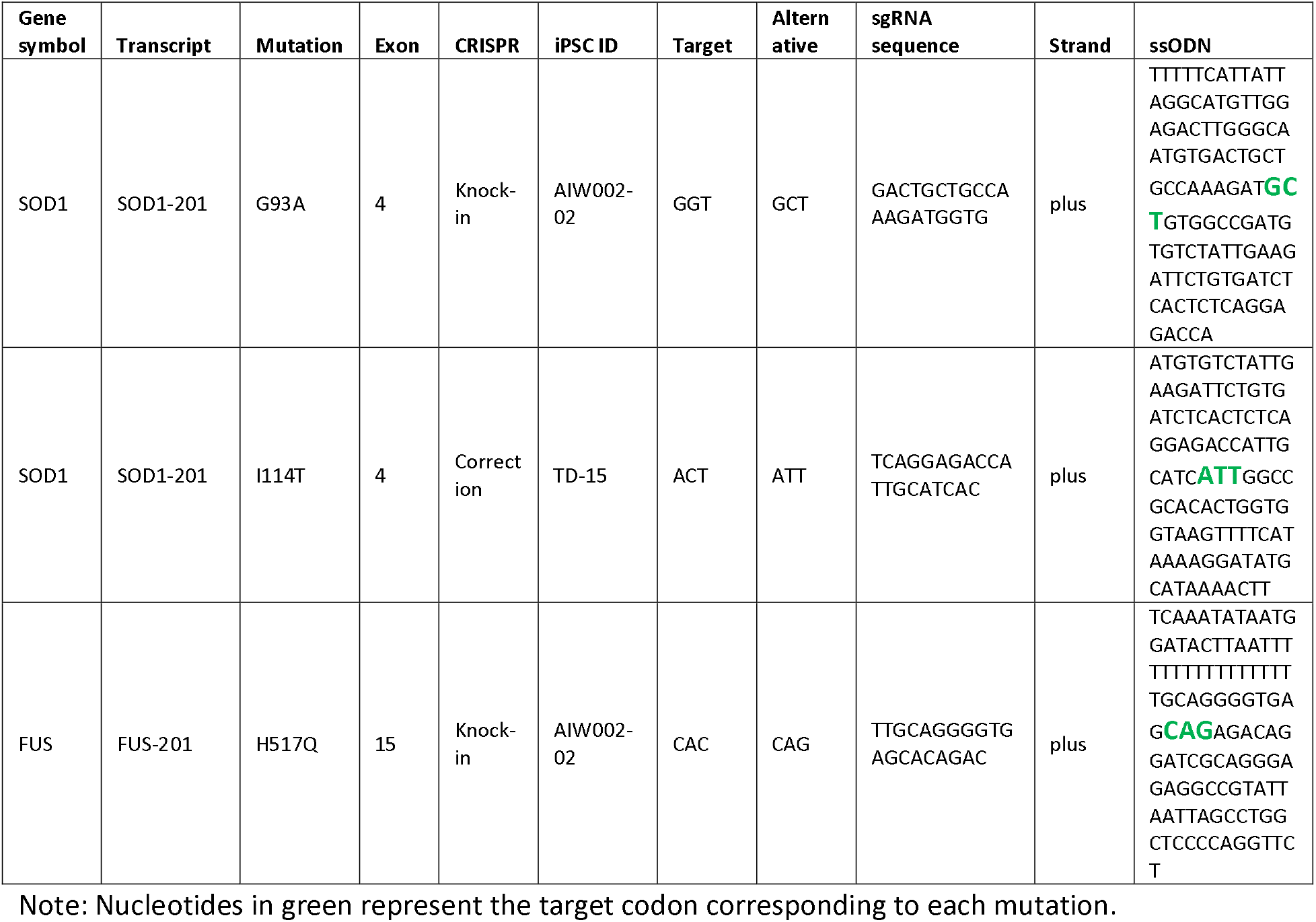
Identification of target codons and design of sgRNAs and ssODNs.

### 3.2. Detection and enrichment of edited alleles

Since HDR typically occurs at low frequency [17], we used ddPCR for accurate quantification of cells containing the desired editing event within a population. For each target, nucleofected iPSCs were uniformly segregated into a 96-well culture plate, allowing the repopulation of each well by a limited cell number in order to enrich for edited cells. The proportion of properly edited alleles was measured in each well using ddPCR with primer/probe sets designed to discriminate between mutant versus corrected (or wt) target codons. We used locked nucleic acid (LNA) probes, with bridged nucleotides at specific positions, to significantly improve affinity and 1-bp-mismatch discrimination [18]. For example, the probe recognizing the G93A mutant allele in *SOD1* contained 6 locked nucleotides in total, including 3 consecutive locked nucleotides encompassing the corresponding point mutation (**Table 5**; see Methods section for details on other stringent design criteria).

**Table 5.**
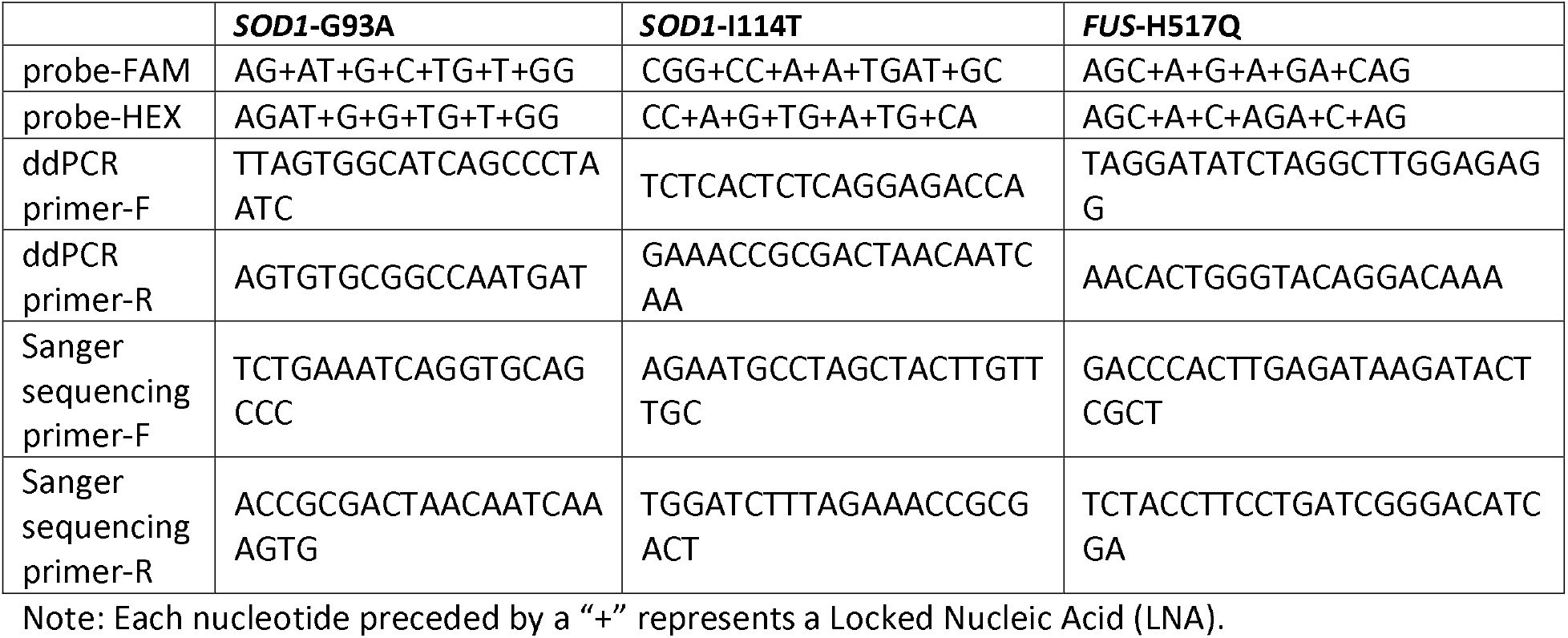
Sequence of primers and probes used for ddPCR or Sanger sequencing.

### 3.3. Correction of I114T in SOD1

We first sought to correct the *SOD1*-I114T missense mutation in our patient iPSC line *SOD1*^I114T/+^. We ensured that the ddPCR-LNA method explained above could efficiently reveal the heterozygous status in our patient line with respect to the I114T missense mutation in *SOD1* (**Fig. 1B**). Applying the same method to the whole 96-well plate post-nucleofection and calculating the proportion of corrected (or wt) alleles led to the identification of seven different wells containing 100% corrected (or wt) alleles (highlighted in yellow in **Fig. 1C**), meaning that no mutant *SOD1*^I114T^ allele was detectable in these wells. These seven different fully-edited cell populations were next monitored for the zygosity of corrected (or wt) alleles to avoid situations where one allele would have acquired an unintended NHEJ-induced insertion-deletion (indel) large enough to impede ddPCR amplification and/or probe recognition. For this, we replaced the I114T mutant probe with an additional primer/probe set in order to co-amplify and detect an unrelated endogenous autosomal gene, i.e., *USP19*, within the same reaction mix. The calculated ratio of *SOD1*^+^:*USP19*^+^ alleles was 1:1 in all of the seven selected wells (**Fig. 1D**), confirming the homozygous status (*SOD1*^+/+^) of the seven cell populations. We chose the iPSCs in well H02 to further perform supporting experiments. The ddPCR scatter plot from the analysis in **Figure 1C** revealed that the mutant alleles previously observed with our patient line (**Fig. 1B**) had completely disappeared (**Fig. 1E**). PCR cloning with external primers and Sanger sequencing performed on the genomic DNA extracted from iPSCs in well H02 confirmed the expected editing event, i.e., the exclusive presence of the codon “ATT” at position 114, without unintended indels (**Fig. 1F**). We verified by short tandem repeat (STR) analysis that the iPSCs in well H02 truly originated from the patient iPSC line and not from any other contaminating *SOD1*^+/+^ cell line (**Fig. 1G**). No further limiting-dilution enrichment steps were necessary for correction of *SOD1*-I114T to generate the isogenic and patient pair.

### 3.4. Introduction of G93A in SOD1

As opposed to correction of *SOD1*-I114T in iPSCs from a heterozygous patient, we next planned to introduce the *SOD1*-G93A mutation into our control iPSC line AIW002-02. We applied the equivalent ddPCR-LNA method explained above for detection of the G93A mutation and determined that our control iPSC line was devoid of *SOD1*^G93A^ alleles (**Fig. 2A**). In the 96-well plate post-nucleofection, we identified 10 different wells presenting cells with 100% mutant alleles (**Fig. 2B**). However, only three of these wells included cells that survived the preparation process for ddPCR (highlighted in yellow in **Fig. 2B**). The zygosity assessment with the endogenous control *USP19* revealed that all of the three wells included a ratio of *SOD1*^G93A^:*USP19*^+^ alleles of 1:1, confirming the homozygous status (*SOD1*^G93A/G93A^) of these three different cell populations (**Fig. 2C**). Well B06 was chosen to visualize the absence of *SOD1*^+^ alleles (**Fig. 2D**), as previously detected in **Figure 2B**. PCR cloning with external primers followed by Sanger sequencing confirmed that only the mutant codon “GCT” was detectable at position 93 without unintended indels (**Fig. 2E**). As for SOD1-I114T correction above, no further limiting-dilution enrichment steps were necessary for the homozygous knock-in of *SOD1*-G93A into the control line, and STR analysis confirmed the genotypic identity. The cells from well B06 were expanded for our differentiation protocol aimed at generating MNs (see below).

**Fig. 2.**
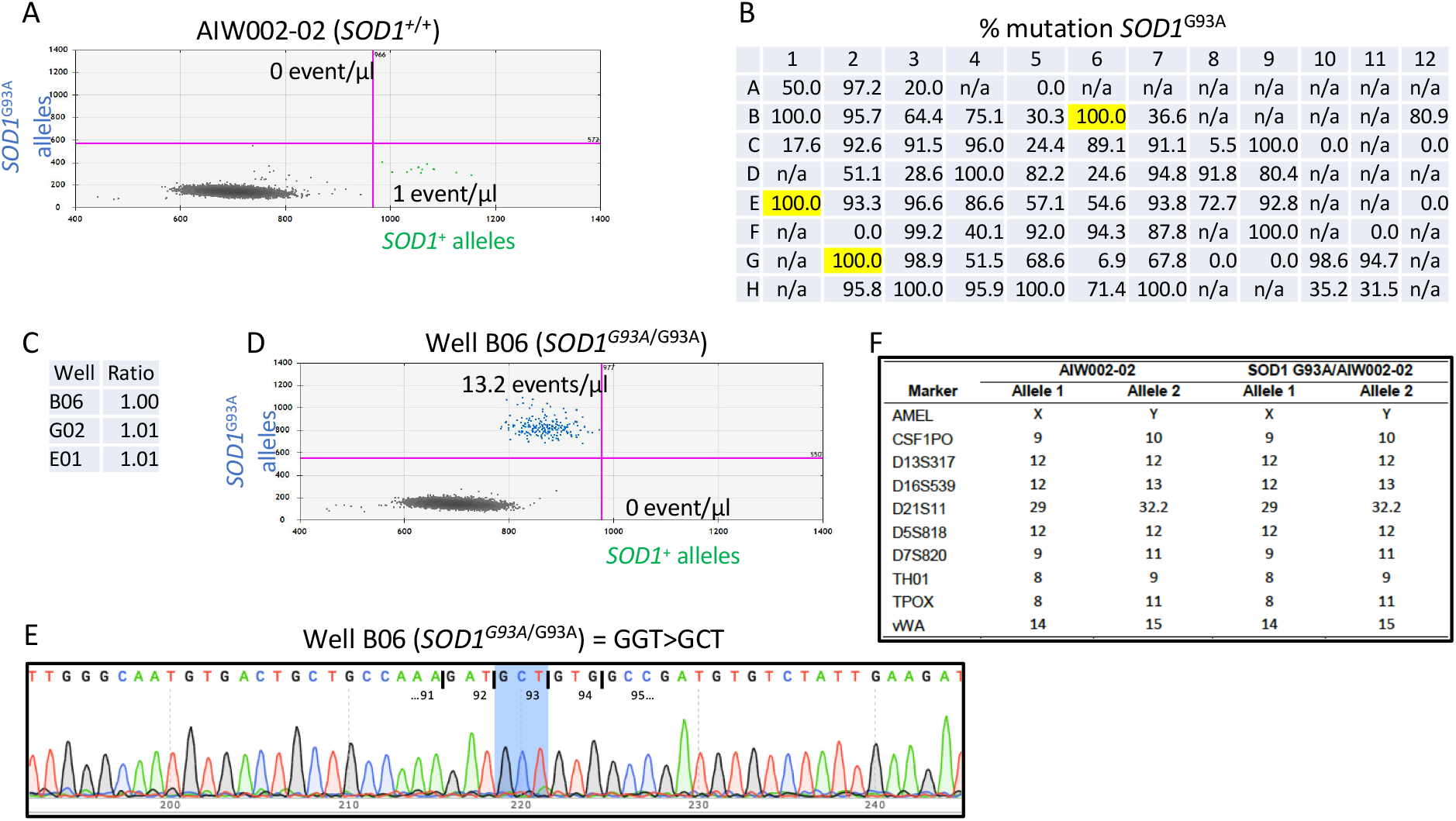
Knock-in of the G93A mutation in *SOD1* in the noncarrier iPSC line AIW002-02. (A) Scatter plot showing the distribution of mutant (blue) vs wt (green) alleles in iPSC line AIW002-02. (B) Percentages of *SOD1*^G93A^ alleles in each well of a 96-well plate measured by ddPCR in the absolute quantification mode. Wells selected for zygosity assessment are highlighted in yellow. (C) Zygosity of the *SOD1*^G93A^ mutation in iPSCs from wells selected in (B), presented as the ratio of *SOD1*^G93A^:*USP19*^+^ alleles. (D) Scatter plot showing the distribution of mutant (blue) vs wt (green) alleles in iPSCs from the well B06 selected in (B). (E) Sanger sequence chromatogram from iPSCs in the selected well B06; shaded blue area shows complete mutation of target codon. (F) Short tandem repeat analysis of line AIW002-02 and cells from well B06. n/a = no colony grew in this well

### 3.5. Introduction of H517Q in FUS

The third CRISPR editing strategy involved the knock-in of the H517Q variant in *FUS* into our control line AIW002-02. The same ddPCR-LNA protocol was used to find only *FUS*^+^ alleles in these iPSCs (**Fig. 3A**). More than 30 different wells containing iPSCs with 100% *FUS*^H517Q^ alleles were defined (**Fig. 3B**). Moreover, all of the wells with live cells had >96% mutant alleles (**Fig. 3B**), suggesting a dramatically high efficiency of editing. However, we were concerned by the possibility of a high rate of NHEJ-induced indel formation preventing ddPCR and allele detection, leaving actually a low rate of true *FUS*^H517Q^ alleles. For example, what appears to be 100% *FUS*^H517Q^ alleles could in fact be only 20% *FUS*^H517Q^ alleles and 80% undetectable alleles bearing indels that preclude ddPCR. Such indels could be due to the fact that the PAM sequence had not been disrupted and no other silent mutation was introduced in the ssODN template (**Table 4**), allowing Cas9 to cut the newly engineered alleles indefinitely until the formation of NHEJ-induced indels interfered with ddPCR and/or probe recognition. As expected, the zygosity profiling for the ratio *FUS*^H517Q^:*USP19*^+^ alleles performed on eight randomly selected wells showed ratios between 0.54 and 0.71 (**Fig. 3C**). Such ratio below 1.0 and the fact that no wt allele was detected, for example in well D07 (**Fig. 3B and D**), suggest the presence of indels that make some alleles undetectable using our ddPCR assay. However, a ratio >0.5 theoretically implies that some cells within the population were homozygous for H517Q and could be isolated upon further limiting-dilution enrichment steps. Well D07, with a ratio of 0.68, was chosen to perform further enrichment of homozygous mutant cells by limited-dilution cell passaging in a secondary 96-well plate, as described [17, 19]. The zygosity test on the secondary plate revealed a ratio 1:1 in well B10 (**Fig. 3E**), indicating that this well was populated only by *FUS^H517Q/H517Q^* iPSCs. In support of this, the scatter plot profile of well B10 displays equivalent numbers of *FUS*^H517Q^ and *USP19*^+^ alleles (**Fig. 3F**). PCR cloning with external primers and Sanger sequencing of DNA from well B10 confirmed the exclusive presence of the codon “CAG” at position 517, without unintended indels (**Fig. 3G**). STR analysis confirmed the genotypic concordance between cells from well B10 and our control line AIW002-02.

**Fig. 3.**
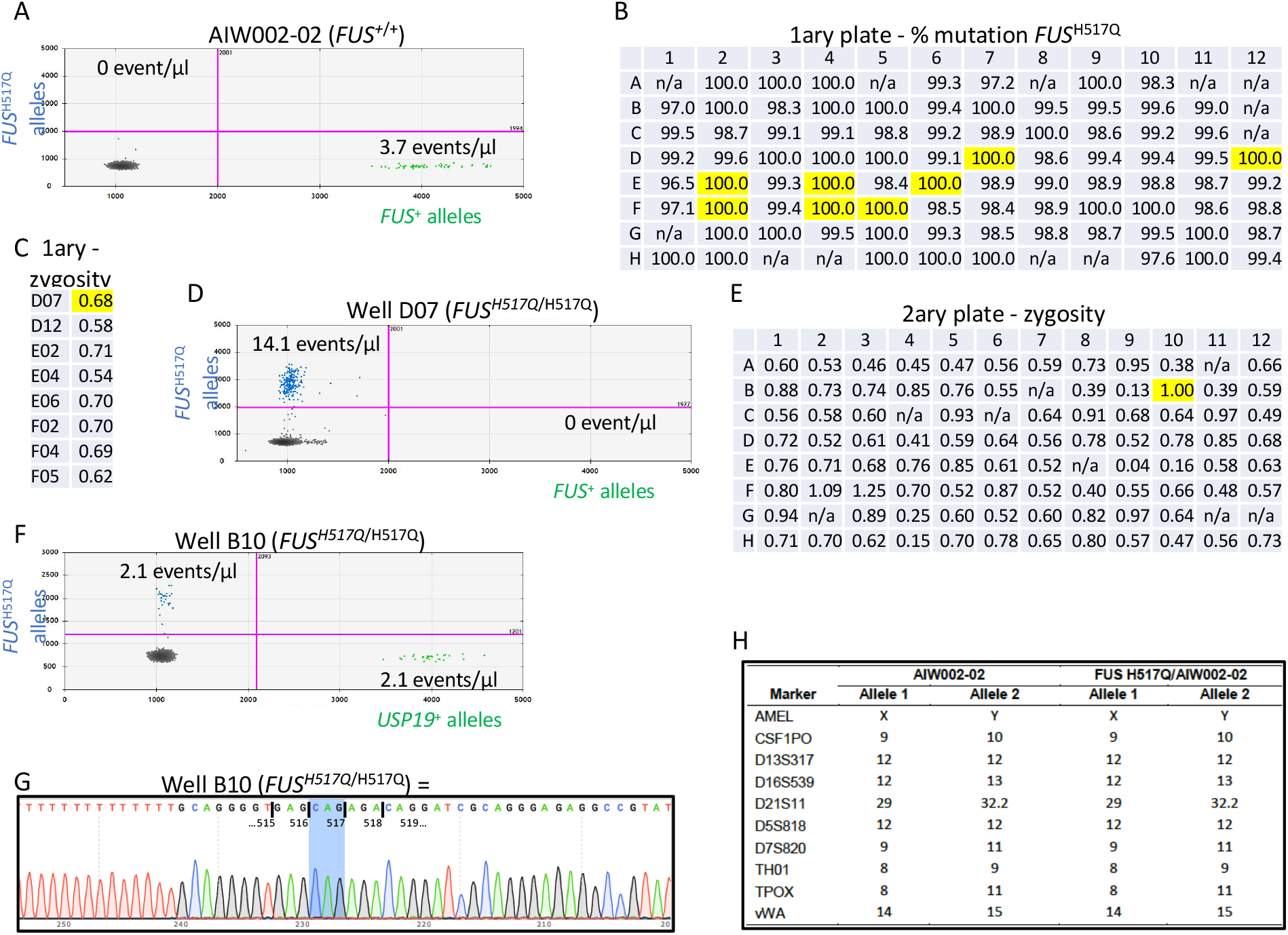
Knock-in of the H517Q mutation in *FUS* in the noncarrier iPSC line AIW002-02. (A) Distribution of mutant (blue) vs wt (green) alleles in iPSC line AIW002-02. (B) Proportion of mutant *FUS*^H517Q^ alleles in the primary 96-well plate measured by ddPCR. Wells selected for zygosity assessment are highlighted in yellow. (C) Zygosity of the *FUS*^H517Q^ mutation in iPSCs from wells selected in (B), presented as the ratio mutant *FUS*^H517Q^:*USP19*^+^ alleles. (D) Scatter plot showing the distribution of mutant (blue) vs wt (green) alleles in iPSCs from well D07 selected in (B). (E) Zygosity of the *FUS*^H517Q^ mutation in each well of the secondary 96-well plate, i.e., limiting-dilution of well D07 from primary plate, measured by ddPCR in the absolute quantification mode, and presented as the ratio of *FUS*^H517Q^:*USP19*^+^ alleles. (F) Scatter plot showing the distribution of mutant (blue) vs wt (green) alleles in iPSCs from the well B10 selected in (**E**). (**G**) Sanger sequence chromatogram from iPSCs in the selected well BIO; shaded blue area shows complete mutation of target codon. (**H**) Short tandem repeat analysis of line AIW002-02 and cells from well B10. n/a = no colony grew in this well

### 3.6. In vitro differentiation and molecular characterization of control and ALS SOD1-G93A motor neurons

Next, to demonstrate that these edited lines could be applied towards generating disease relevant cells, we differentiated MNs from either our control iPSCs or iPSCs carrying the ALS risk variant G93A using a small molecule-based approach that recapitulates the developmental process of spinal cord MNs *in vivo* [16]. Briefly, neural induction and caudalization of the neural progenitor cells was achieved through activation of WNT signalling along with the inhibition of BMP/TGFbeta signaling with the small molecules CHIR99021, SB431542 and DMH1. To promote MN differentiation, the subsequent ventralization and caudalization of the NPCs was realized through the addition of Purmorphamine, to activate shh signalling, and retinoic acid (RA), respectively. Finally, to prevent neurogenesis and to keep cells at the NPC stage before starting the final differentiation into MNs, valproic acid (VPA) was added to the media (**Fig. 4A**). iPSCs, MN NPCs and MNs differentiated for 14 and 28 days were analysed by qPCR and an additional characterization of the MN NPCs and MNs following their differentiation for 28 days was done by immunocytochemistry (ICC) for both the control line AIW002-02 and the knock-in line G93A. iPSCs from both lines expressed the pluripotency markers NANOG and OCT4 and no expression of these markers was detected either in the MN NPCs or in the differentiated MNs, showing that no iPSCs remained in these cultures over the course of differentiation (**Fig. 4D**). The AIW002-02 and SOD1-G93A iPSC lines generated neural progenitor cells (NPCs) expressing the NPC markers Sox1 and Nestin (**Fig. 4B**). These NPCs were proliferative, as demonstrated by the expression of the cell proliferation marker Ki67 by ICC (**Fig. 4B**). Additionally, following the ventralization and caudalization of the NPCs by the addition of Purmorphamine and RA, the specification of the NPCs into MN NPCs was confirmed by the expression of Pax6 and the MN NPC marker Olig2 in both lines (**Fig. 4D**). Expression of the neuronal markers MAP2 and NF-H was confirmed by qPCR (**Fig. 4D**) and ICC (**Fig. 4C**) analysis respectively, in MNs at 14 and 28 days of differentiation, showing the efficacy of the neuronal differentiation for both lines. Additionally, these neurons expressed the MN markers HB9 and Islet1 as early as 14 days of differentiation (**Fig. 4C and D**). The MN identity obtained was further confirmed by the co-expression of the choline acetyltransferase protein ChAT along with Hb9 and Islet1 (**Fig. 4C and D**). Together these observations provide evidence for the generation of neurons presenting a molecular profile typical of MNs, from both the human iPSC lines knock-in ALS *SOD1* mutant G93A, and its isogenic control AIW002-02.

**Fig. 4.**
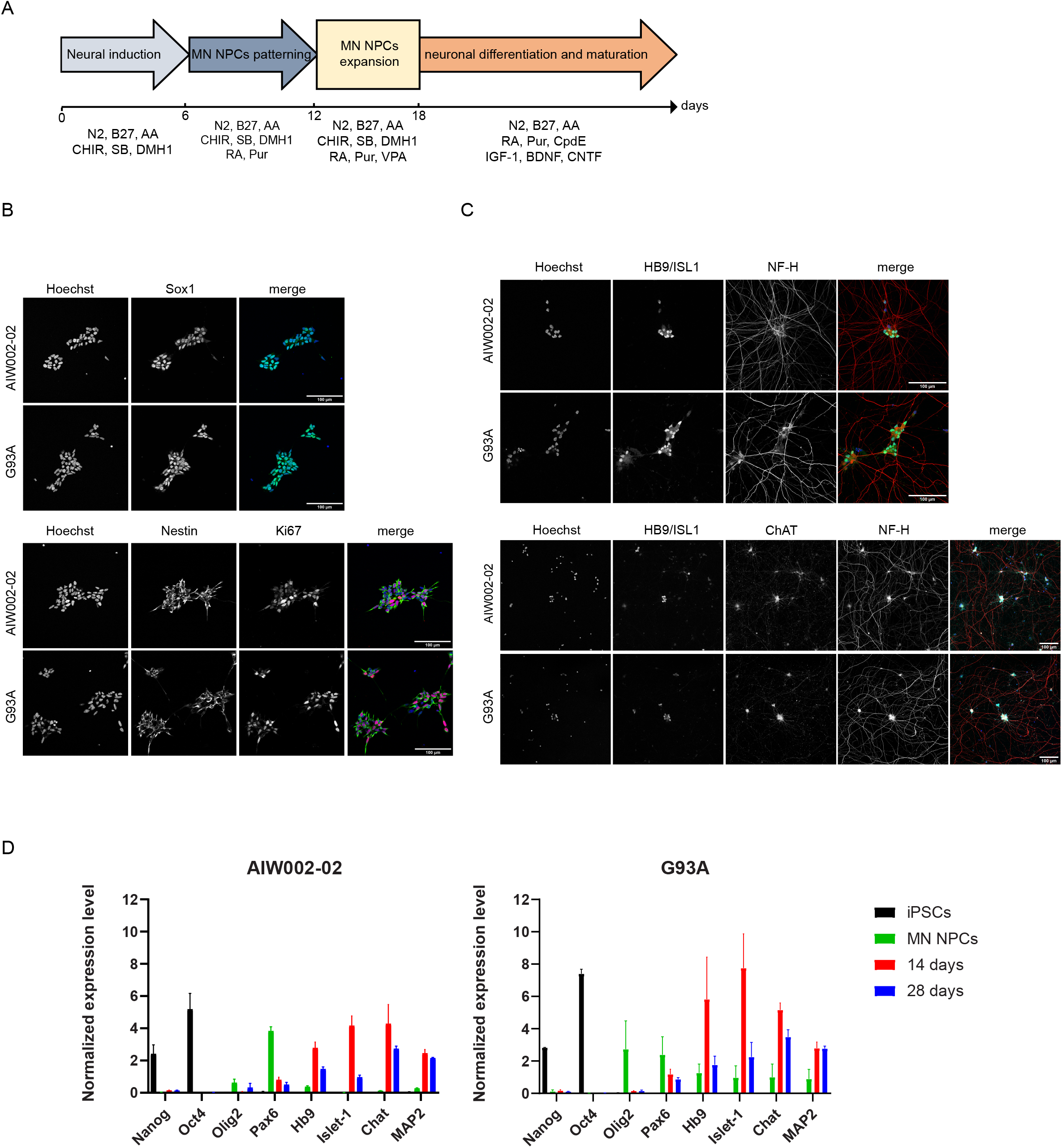
Differentiation of human iPSCs into motor neurons. (A) Schematic representation of the *in vitro* differentiation protocol used to generate iPSC-derived MNs. The time-course and combination of small molecules used for each step is shown. AA: Ascorbic Acid, CHIR: CHIR99021, SB: SB431542, RA: Retinoic Acid, Pur: Purmorphamine, VPA: Valproic Acid, IGF-1: Insulin Growth Factor 1, BDNF: Brain derived neurotrophic factor, CNTF: Ciliary Neurotrophic Factor. (B) Representative images of AIW002-02 and *SOD1*^G93A^ iPSC-derived NPCs visualized by immunocytochemistry. Cells were stained with the neural progenitor markers Sox1 and Nestin, and the proliferative marker Ki67. Nuclei were counterstained with Hoechst 33342. (C) Representative images of AIW002-02 and *SOD1*^G93A^ iPSC-derived MNs visualized by immunocytochemistry with the combined HB9 and Islet1 MN markers (HB9/ISL1), the choline acetyltransferase protein (ChAT), and the neuronal marker neurofilament heavy chain (NF-H) after 28 days of final differentiation. Nuclei were counterstained with Hoechst 33342. (D) qPCR depicting normalized expression levels of pluripotency markers NANOG and OCT4, MN NPCs markers OLIG2 and Pax6, MNs markers HB9, ISL1 and CHAT, and the neuronal marker MAP2, in iPSCs, MN NPCs and D14 and D28 iPSC-derived motor neurons from the *SOD1* G93A knock-in and its isogenic control AIW002-02. Data were normalized to Actβ-GAPDH expression. Bar graphs show mean +/-SEM; n = 3 independent experiments (except from G93A knock-in iPSCs where n=2), with each data point obtained from a separate culture.

## 4. Discussion

Neurological disorders such as ALS can be studied using patient-derived iPSCs, in which candidate risk factors with variable penetrance can be modeled and manipulated using CRISPR gene editing. However, small and precise DNA modifications such as point mutations are not always straightforward to reproduce. Here, we present a quick, easy and efficient CRISPR-Cas9 method to edit single point mutations associated with ALS in human iPSCs without the use of antibiotic selection (**Fig. 1A**). These mutations were corrected in patient iPSCs, i.e., I114T in *SOD1*, or introduced into control iPSCs, i.e., G93A in *SOD1* and H517Q in *FUS* (**Table 4**). The purpose of such precise genomic DNA modifications is to generate isogenic pairs of MNs for the functional validation of causal DNA variants. Small effect sizes are often observed with genetic variants associated with ALS [2]. Thus, the power to detect significant phenotypic changes is increased by reducing variance through employing isogenic pairs.

New gene editing techniques including CRISPR-Cas9 provide the opportunity to knockout candidate genes to study their function in the etiology of ALS, and also to manipulate individual SNP to study the role of pathologic genetic variants. However, substitution of an individual nucleotide can be particularly challenging in the context of current CRISPR tool availability since it is a rare event, often occurring at <1% frequency. To cope with this hurdle, antibiotic resistance genes are typically used to select for properly-edited cells, but unfortunately leave behind exogenous remnant sequences that can interfere with cellular phenotypes. To avoid antibiotic selection and leftover DNA, we adapted a method using ddPCR where thousands of single-genome PCR reactions are simultaneously performed to permit the absolute quantification of rare editing events within a cell population, coupled with limiting-dilution selection steps [17, 19, 20]. However, the specificity of the designed Taqman probes used in the ddPCR reaction is often low, to the point where edited and nonedited alleles cannot be efficiently discriminated during the analysis. Hence, we have integrated here the use of LNA probes, with bridges between specific nucleotides to considerably increase specificity and mismatch discrimination [18]. Furthermore, the efficiency of editing was high enough so that no additional limiting-dilution steps were necessary to isolate a 100%-edited clone for two of our three target genes.

Potential indels interfering with the ddPCR reaction were found in a considerable number of wells in the primary plate for the *FUS*^H517Q^ knock-in, but not for *SOD1*^G93A^, which was likely because of the absence of a silent mutation in the *FUS*^H517Q^ ssODN template to disrupt the PAM sequence. Through this approach, Cas9 was likely able to cut the newly engineered alleles multiple times until the appearance of an indel prevented it from cutting any further. However, the substitution of the single nucleotide necessary for the correction of *SOD1*^G93A^ was sufficient to disrupt the PAM sequence. For the correction of *SOD1*^I114T^ in our patient line, we hypothesize that the sgRNA-Cas9 molecule could not cut the wt allele since the SNP was only 1 bp away from the PAM, precluding the formation of any indels on the original wt allele.

Various types of neurons involved in diverse neurological disorders may be generated from iPSCs by exposure to different combinations of small molecules and developmental morphogens [16, 21, 22]. Alternatively, rapid and highly efficient neuronal differentiation can also be achieved through forced expression of defined transcription factors [23–25]. In this work, we used an approach based on small molecules to differentiate isogenic iPSCs so as to reflect as close as possible the *in vivo* developmental trajectory of spinal cord MNs [16]. Using the example of the SOD1-G93A knocked-in mutation and its isogenic control AIW002-02 line, we demonstrated our ability to differentiate our CRISPR-edited iPSCs into MNs as efficiently as its isogenic control. Having the possibility to introduce mutations of interest not available in patients, or to correct mutations in order to have access to isogenic controls for some patient’s lines will help us to identify potential ALS phenotypic variations with specific genetic models of ALS. We believe that the various iPSCs engineered in this work will serve as models of ALS to generate further insights into the fundamental mechanisms underlying neuron degeneration in ALS.

## CRediT authorship contribution statement

**Eric Deneault:** Conceptualization, Data curation, Formal analysis, Investigation, Methodology, Writing - original draft. **Mathilde Chaineau:** Conceptualization, Data curation, Formal analysis, Methodology, Writing - original draft. **Maria José Castellanos Montiel, Anna Krystina Franco Flores, Ghazal Haghi, Carol XQ Chen, Narges Abdian, Michael Nicouleau:** Data curation, Formal analysis, Methodology. **Irina Shlaifer:** Formal analysis, Methodology, Writing - original draft. **Lenore K. Beitel:** Writing - review & editing. **Thomas M. Durcan:** Conceptualization, Funding acquisition, Project administration, Resources, Supervision, Validation, Writing - review & editing.

## Acknowledgements

TMD received funding to support this project from the Canada First Research Excellence Fund, awarded through the Healthy Brains, Healthy Lives initiative at McGill University, the CQDM FACS research program and a US DOD ALS research grant. TMD is supported by a project grant from CIHR (PJT–169095). We wish to thank the C-BIG Biorepository Histology Core Facility (C-BIG) for cell repository, and Geneviève Dorval for technical help.

